# Deterministic splicing of *Dscam2* is regulated by Muscleblind

**DOI:** 10.1101/297101

**Authors:** Joshua Shing Shun Li, S.Sean Millard

## Abstract

Alternative splicing of genes increases the number of distinct proteins in a cell. In the brain it is highly prevalent, presumably because proteome diversity is crucial for establishing the complex circuitry between trillions of neurons. To provide individual cells with different repertoires of protein isoforms, however, this process must be regulated. Previously, we found that the mutually exclusive alternative splicing of a cell surface protein, *Dscam2* produces two isoforms (exon 10A and 10B) with unique binding properties. This splicing event is tightly regulated and crucial for maintaining axon terminal size, dendritic morphology and synaptic numbers. Here, we show that *Drosophila* Muscleblind (Mbl), a conserved splicing factor implicated in myotonic dystrophy, controls *Dscam2* alternative splicing. Removing *mbl* from cells that normally express isoform B induces the expression of isoform A and eliminates the expression of B, demonstrating that Mbl represses one alternative exon and selects the other. *Mbl* mutants exhibit phenotypes that are also observed in flies engineered to express a single isoform. Consistent with these observations, *mbl* expression is cell-type-specific and correlates with the expression of isoform B. Our study demonstrates how the regulated expression of a splicing factor is sufficient to provide neurons with unique protein isoforms crucial for development.

## Introduction

Alternative splicing occurs in approximately 95% of human genes and generates proteome diversity much needed for brain wiring (Pan et al., 2008; Wang et al., 2008). Specifying neuronal connections through alternative splicing would require regulated expression of isoforms with unique functions in different cell types to carry out distinct processes. Although there are some examples of neuronal cell-type-specific isoform expression (Bell et al., 2004; Iijima et al., 2014; Lah et al., 2014; Norris et al., 2014; Schreiner et al., 2014; Tomioka et al., 2016), the mechanisms underlying these deterministic splicing events and their functional consequences remain understudied. This is due, in part, to the technical difficulties of assessing and manipulating isoform expression *in vivo*, and at the single cell level. Another obstacle is that most splicing regulators are proposed to be ubiquitously expressed (Nilsen and Graveley, 2010). For example, the broadly expressed SR and heterogeneous nuclear ribonucleoproteins (hnRNPs) typically have opposing activities, and the prevalence of splice site usage is thought to be controlled by their relative abundances within the cell (Blanchette et al., 2009). Although there are many examples where splicing regulators are expressed in a tissue-specific manner (Calarco et al., 2009; Kuroyanagi et al., 2006; Markovtsov et al., 2000; Ohno et al., 2008; Underwood et al., 2005; Warzecha et al., 2009), until recently, reports of cell-type specific expression have been less frequent (McKee et al., 2005; Wang et al., 2018).

In insects, Dscam2 is a cell recognition molecule that mediates self- and cell-type-specific avoidance (tiling) (Funada et al., 2007; Millard et al., 2007; Millard et al., 2010). Mutually exclusive alternative splicing of exon 10A or 10B produces two isoforms with biochemically unique extracellular domains that are regulated both spatially and temporally (Funada et al., 2007; Millard et al., 2007). Previously, we demonstrated that cell-type-specific alternative splicing of *Drosophila Dscam2* is crucial for the proper development of axon terminal size, dendrite morphology and synaptic numbers in the fly visual system (Kerwin et al., 2018; Lah et al., 2014; Li et al., 2015). Although these studies showed that disrupting cell-specific *Dscam2* alternative splicing has functional consequences, what regulates this process remained unclear. Here, we conducted an RNAi screen and identified *muscleblind* (*mbl*) as a regulator of *Dscam2* alternative splicing. Loss-of-function (LOF) and overexpression (OE) studies suggest that Mbl acts both as a splicing repressor of *Dscam2* exon 10A and as an activator of exon 10B (hereafter *Dscam2.10A* and *Dscam2.10B*). Consistent with this finding, *mbl* expression is cell-type-specific and correlates with the expression of *Dscam2.10B*. Hypomorphic *mbl* mutants exhibit visual system phenotypes that are similar to those observed in flies engineered to express one isoform in all *Dscam2*-positive cells (single isoform strains). Similarly, driving *mbl* in mushroom body neurons that normally select isoform A, induces the expression of isoform B and generates a single isoform phenotype. Although the *mbl* gene is itself alternatively spliced, we found that selection of *Dscam2.10B* does not require a specific Mbl isoform and that human MBNL1 can also regulate *Dscam2* alternative splicing. Our study provides compelling genetic evidence that the regulated expression of a highly conserved RNA binding protein, Mbl, is sufficient for the selection of *Dscam2.10B* and that disrupting this mechanism for cell-specific protein expression leads to developmental defects in neurons.

## Results

### An RNAi screen identifies *mbl* as a repressor of *Dscam2* exon 10A selection

We reasoned that the neuronal cell-type-specific alternative splicing of *Dscam2* is likely regulated by RNA binding proteins, and that we could identify these regulators by knocking them down in a genetic background containing an isoform reporter. In photoreceptors (R cells) of third instar larvae, *Dscam2.10B* is selected whereas the splicing of *Dscam2.10A* is repressed (Lah et al., 2014; Tadros et al., 2016). Given that quantifying a reduction in *Dscam2.10B* isoform reporter levels is challenging compared to detecting the appearance of *Dscam2.10A* in cells where it is not normally expressed, we performed a screen for repressors of isoform A in R cells.

To knock down RNA binding proteins, the *glass* multimer reporter (*GMR)-GAL4* was used to drive RNAi transgenes selectively in R cells. Our genetic background included *UAS-Dcr-2* to increase RNAi efficiency (Dietzl et al., 2007) and *GMR-GFP* to mark the photoreceptors independent of the *Gal4/UAS* system (Brand and Perrimon, 1993). Lastly, a *Dscam2.10A-LexA* reporter driving *LexAOp*-myristolated tdTomato (hereafter *Dscam2.10A>tdTom*; Fig. 1A) was used to visualize isoform A expression (Lai and Lee, 2006; Tadros et al., 2016). As expected, *Dscam2.10B>tdTom* was detected in R cell projections in the lamina plexus as well as in their cell bodies in the eye-disc, whereas *Dscam2.10A>tdTom* was not (Fig. 1C-1D). Overexpression of Dcr-2 in R cells did not perturb the repression of *Dscam2.10A* (Fig 1O). We knocked down ~160 genes using ~250 RNAi lines (Fig 1B and Table S1) and identified two independent RNAi lines targeting *mbl* that caused aberrant expression of *Dscam2.10A* in R cells where it is normally absent (Fig 1F, 1O). The penetrance increased when animals were reared at a more optimal Gal4 temperature of 29°C (Mondal et al., 2007; Ni et al., 2008) (Fig 1O).

**Figure 1.**
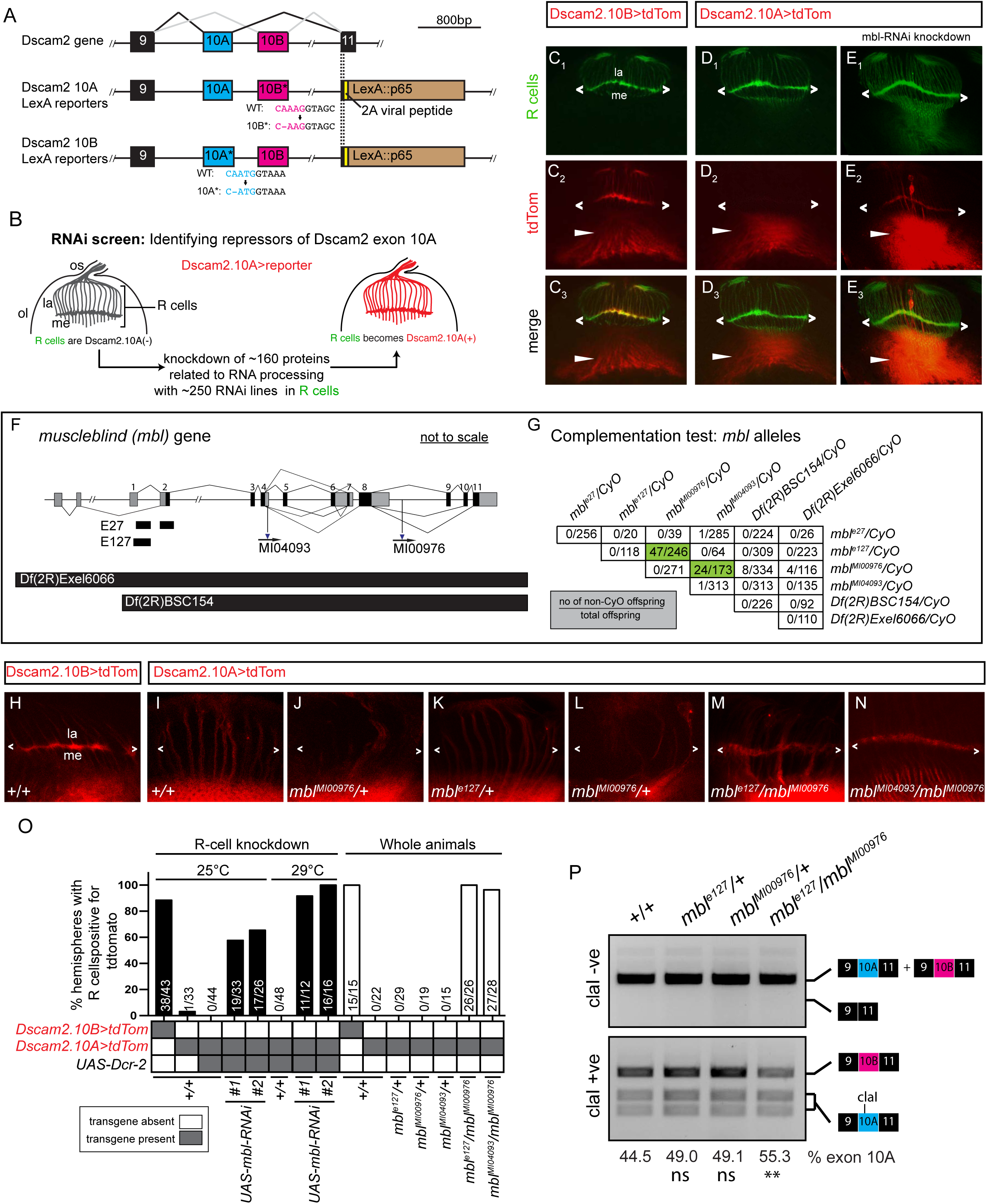
*Drosophila mbl* is required for the repression of *Dscam2* exon 10A in R cells. (A) Schematic showing the region of *Dscam2* exon 10 that undergoes mutually exclusive alternative splicing and the LexA isoform-specific reporter lines. Frame-shift mutations in the exon not reported are shown. (B) Schematic RNAi screen design for identifying repressors of *Dscam2* exon 10A selection. R cells normally select exon 10B and repress exon 10A. We knocked-down RNA binding proteins in R cells while monitoring 10A expression.(C-E) *Dscam2* exon 10A is derepressed in R cells when *mbl* is knocked-down. (C_1_-C_3_) *Dscam2.10B* control. R cells (green) normally select exon 10B (red). R cell terminals can be observed in the lamina plexus (angle brackets). *Dscam2.10B* is also expressed in the developing optic lobe (arrowhead). (D_1_-D_3_) *Dscam2.10A* is not expressed in R cells (green) but is expressed in the developing optic lobe (arrowhead). (E_1_-E_3_) RNAi lines targeting *mbl* in R cells results in the aberrant expression of *Dscam2.10A* in R cells. (F) Schematic of the *mbl* gene showing the location of two small deletions (*E27* and *E127*), two MiMIC insertions (*MI04093* and *MI00976*) and two deficiencies (*Df(2R)Exel6066 and Df(2R)BSC154*) used in this study. Non-coding exons are in gray, coding exons are black.(G) Complementation test of *mbl* loss-of-function (LOF) alleles. Numbers in the table represent the number of non-*CyO* offspring over the total. Most transheterygote combinations were lethal with the exception of *mbl*^*MI00976*^*/mbl*^*e27*^and *mbl*^*MI00976*^*/mbl*^*MI04093*^(green). (H-N) *Mbl* transheterozygotes express *Dscam2.10A* in R cells. (H) *Dscam2.10B* control showing expression in the lamina plexus (angle brackets). (I) *Dscam2.10A* control showing no expression of this isoform in R cells. (J-L) Heterozygous animals for *mbl* LOF alleles are comparable to control. (M-N) Two different *mbl* transheterozygote combinations exhibit de-repression of *Dscam2.10A* in R cells. (O) Quantification of *Dscam2.10>tdTom* expression in third instar R cells with various *mbl* manipulations; including RNAi knockdown (black bars) and whole animal transheterozygotes (white). Y-axis represents the number of optic lobes with R cells positive for tdTom over total quantified as a percentage. On the x-axis, the presence of a transgene is indicated with a grey box and the temperature at which the crosses were reared (25°C or 29°C) is indicated on the top. (P) *Dscam2* exon 10A inclusion is increased in *mbl* transheterozygotes. (Top) Semiquantitative RT-PCR from different genotypes indicated. Primers amplified the variable region that includes exon 10. A smaller product that would result from exon 10 skipping is not observed. (Bottom) Exon 10A-specific cleavage with restriction enzyme ClaI shows an increase in exon 10A inclusion in *mbl* transheterozygotes. Percentage of exon 10A inclusion was calculated by dividing 10A by 10A+10B bands following restriction digest. The mean of exon 10A inclusion is shown at the bottom of each lane. ANOVA test with Tukey’s multiple comparison test was used to compare the exon 10A inclusion. ns *P* > 0.05, ** *P* < 0.01. See also Figures S1 and S2.

Mbl-family proteins possess evolutionarily conserved tandem CCCH zinc-finger domains through which they bind pre-mRNA. Vertebrate Mbl family members are involved in tissue-specific splicing and have been implicated in myotonic dystrophy (Pascual et al., 2006). Formerly known as *mindmelt, Drosophila mbl* was first identified in a second chromosome *P*-element genetic screen for embryonic defects in the peripheral nervous system (Kania et al., 1995). *Mbl* produces multiple isoforms through alternative splicing (Begemann et al., 1997; Irion, 2012), and its function has been most extensively characterized in fly muscles where both hypomorphic mutations and sequestration of the protein by repeated CUG sequences within an mRNA lead to muscle defects (Artero et al., 1998; Llamusi et al., 2013). To validate the RNAi phenotype, we tested *Dscam2.10A>tdTom* expression in *mbl* loss-of-function (LOF) mutants. Since *mbl* LOF results in lethality, we first conducted complementation tests on six *mbl* mutant alleles to identify viable hypomorphic combinations. These included two alleles created previously via imprecise *P*-element excision (*mbl*^*e127*^and *mbl*^*e27*^; Begemann et al. 1997) two MiMIC splicing traps (*mbl*^*MI00976*^and *mbl*^*MI04093*^; (Venken et al., 2011) and two 2^nd^chromosome deficiencies (*Df(2R)BSC154* and *Df(2R)Exel6066*; Fig 1F-1G). Consistent with previous reports, the complementation tests confirmed that the majority of the alleles were lethal over one another (Fig 1G) (Kania et al., 1995). However, we identified two *mbl* transheterozygous combinations that were partially viable and crossed these into a *Dscam2.10A>tdTom* reporter background. Both *mbl*^*e127*^*/mbl*^*MI00976*^and *mbl*^*MI04093*^*/mbl*^*MI00976*^animals presented aberrant *Dscam2.10A* expression in R cells when compared to heterozygous and wild-type controls (Fig 1H-O). *Mbl* mutant mosaic clones also exhibited aberrant *Dscam2.10A>tdTom* expression in R cells (Fig S1A-S1F). The weakest allele, *mbl*^*M00976*^, which removes only a proportion of the *mbl* isoforms, was the only exception (Fig S1E-S1F).

One alternative explanation of how *Dscam2.10A>tdTom* expression could get switched-on in *mbl* mutants, is through exon 10 skipping. Removing both alternative exons simultaneously does not result in a frameshift mutation, and since the Gal4 in our reporters is inserted directly downstream of the variable exons (in exon 11), it would still be expressed. To test this possibility, we amplified *Dscam2* sequences between exon 9 and 11 in *mbl*^*e127*^*/mbl*^*MI00976*^transheterozygous animals using RT-PCR. In both control and *mbl* LOF mutants, we detected RT-PCR products (~690bp) that corresponded to the inclusion of exon 10 (A or B) and failed to detect products (~390bp) that would result from exon 10 skipping (Fig 1P). This suggested that Mbl is not involved in the splicing fidelity of *Dscam2.10* but rather in the selective mutual exclusion of its two isoforms. To assess whether the ratios of the two isoforms were changing in the *mbl* hypomorphic mutants, we cut the exon 10 RT-PCR products with the *ClaI* restriction enzyme that only recognizes exon 10A. Densitometric analysis then allowed us to semi-quantitatively compare the relative levels of both isoforms. There was ~25% increase in the level of exon 10A inclusion in *mbl*^*e127*^*/mbl*^*MI00976*^ animals compared to controls (Fig 1P). Similarly, qRT-PCR of the *mbl*^*e127*^*/mbl*^*MI00976*^animals showed a ~1.25 fold and ~0.78 fold change in exon 10A and 10B inclusion respectively, when compared to controls. Both results are consistent with the de-repression we observed in our 10A reporter lines. To determine whether Mbl was specifically regulating *Dscam2* exon 10 mutually exclusive splicing, we assessed other *Dscam2* alternative splicing events. These included an alternative 5’ splice site selection of *Dscam2* exon 19 and the alternative last exon (ALE) selection of exon 20 (Fig S2A). The expression of these different isoforms was unchanged in *mbl* hypomorphic mutants (Fig S2B). Together, our results indicate that Mbl is an essential splicing factor that specifically represses *Dscam2.10A*.

### Mbl is necessary for the selection of *Dscam2* exon 10B

Since *Dscam2* exon 10 isoforms are mutually exclusively spliced, we predicted that selection of exon 10A would lead to the loss of exon 10B selection. To test this, we conducted mosaic analysis with a repressible cell marker (MARCM) (Lee and Luo, 1999) to analyse *Dscam2.10B* expression in *mbl* mutant clones. In late third instar brains, clones homozygous (GFP-positive) for *mbl*^*e127*^and *mbl*^*e27*^exhibited a dramatic reduction in *Dscam2.10B>tdTom* expression in R cell axons projecting to the lamina plexus compared to controls (Fig 2B, C, E). The absence of *Dscam2.10B>tdTom* in *mbl* mutant clones was more striking during pupal stages (Fig 2D), suggesting that perdurance of Mbl could explain the residual signal observed in third instar animals. These results reveal that *mbl* is cell-autonomously required for the selection of the *Dscam2.10B*.

**Figure 2.**
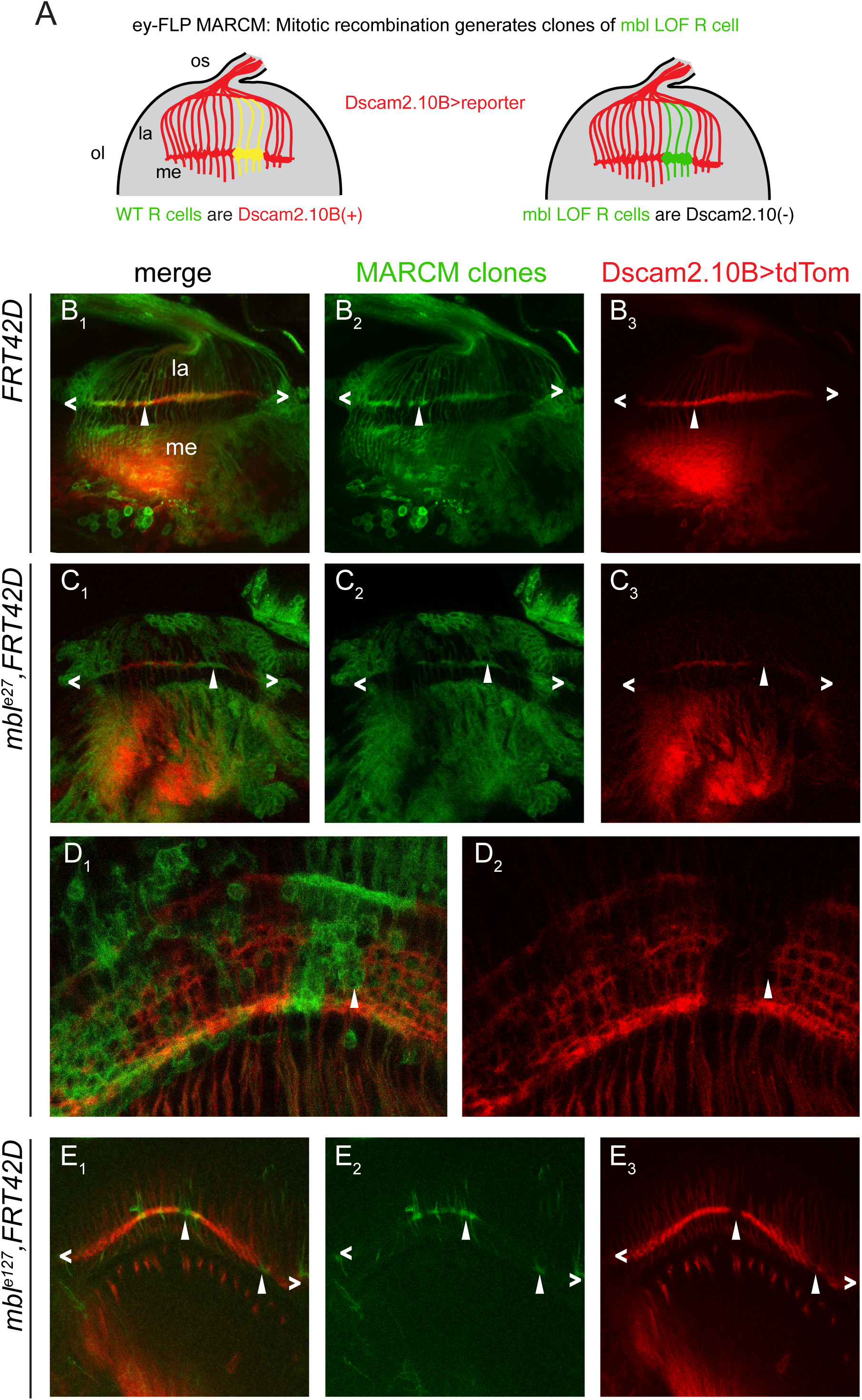
*Drosophila mbl* is necessary for the selection of *Dscam2* exon 10B in R cells. (A) Schematic of our predicted *mbl* MARCM results using *ey-FLP.* WT R cell clones will be GFP(+) and *Dscam2.10B>tdTom*(+) (yellow), whereas *mbl* mutant clones will be *Dscam2.10B>tdTom*(-) (green). (B_1_-B_3_) Control MARCM clones (green) in 3^rd^instar R cells (angle brackets) are positive for *Dscam2.10B>tdTom* (arrowhead). (C_1_-C) In *mbl*^*e27*^clones, *Dscam2.10B* labelling in the lamina plexus is discontinuous and its absence correlates with the loss of Mbl (arrowhead). (D_1_-D_2_) *Mbl* MARCM clones from midpupal optic lobes lack *Dscam2.10B*>tdTom. (E_1_-E_3_) A different allele (*mbl*^*e127*^) exhibits a similar phenotype in third instar brains.

### *mbl* expression is cell-type-specific and correlates with *Dscam2.10B* selection

Previous studies have reported that *mbl* is expressed in third instar eye-discs and muscles (Artero et al., 1998; Brouwer et al., 1997). Since *mbl* LOF results in both the selection of *Dscam2.10A* and the loss of *Dscam2.10B*, we predicted that *mbl* expression would correlate with the presence of isoform B. To test this, we characterized several *mbl* reporters (Fig S3A). We analyzed three enhancer trap strains (transcriptional reporters) inserted near the beginning of the *mbl* gene (*mbl*^*k01212*^*-LacZ, mbl*^*NP1161*^*-Gal4* and *mbl*^*NP0420*^*-Gal4*), as well as a splicing trap line generated by the Trojan-mediated conversion of a *mbl* MiMIC (Minos Mediated Integration Cassette) insertion (Fig S2A, *mbl*^*MiMIC00139*^*-Gal4;* (Diao et al., 2015). The splicing trap reporter consists of a splice acceptor site and an in-frame *T2A-Gal4* sequence inserted in an intron between two coding exons. This *Gal4* cassette gets incorporated into *mbl* mRNA during splicing and therefore Gal4 is only present when *mbl* is translated. Consistent with previous studies, and its role in repressing the production of *Dscam2.10A*, all four *mbl* reporters were expressed in the third instar photoreceptors (Fig 3A, S3A-S3D). We next did a more extensive characterization of *mbl* expression by driving nuclear localized GFP (*GFP.nls*) with one transcriptional (*mbl*^*NP0420*^*-Gal4)* and one translational (*mbl*^*MiMIC00139*^*-Gal4)* reporter. In the brain, we found that *mbl* was expressed predominantly in postmitotic neurons with some expression detected in glial cells (Fig SEC-S3H and S3J-S3M). Interestingly, we detected the translational but not the transcriptional reporter in third instar muscles (Fig S3I and S3N). The absence of expression is likely due to the insertion of the *P*-element into a neural-specific enhancer, as previously described (Bargiela et al., 2014). To assess the expression of *mbl* in the five lamina neurons L1-L5, all of which express *Dscam2* (Lah et al., 2014; Tadros et al., 2016), we implemented an intersectional strategy using a *UAS>stop>epitope* reporter (Nern et al., 2015) that is dependent on both *FLP* and *Gal4*. The *FLP* source (*Dac-FLP*) was expressed in lamina neurons and able to remove the transcriptional stop motif in the reporter transgene. The overlap between *mbl-Gal4* and *Dac-FLP* allowed us to visualize *mbl* expression in lamina neurons at single-cell resolution (Fig 3B). As a proof of principle, we first did an intersectional analysis with a pan-neuronal reporter, *elav-Gal4* (Fig 3C_1_). We detected many clones encompassing various neuronal-cell-types including the axons of L1-L5 and R7-R8 (Fig 3C-3D). This confirmed that all lamina neurons could be detected using this strategy. Using *mbl-Gal4* reporters we found that L1, R7 and R8, which expresses *Dscam2.10B*, were the primary neurons labelled. A few L4 cells were also detected, which is consistent with this neuron expressing *Dscam2.10B* early in development and *Dscam2.10A* at later stages (Tadros et al., 2016). To confirm this finding, we dissected the expression of *mbl* in lamina neurons during development. Using the same intersectional strategy, we detected a high number of L4 clones at 48hr apf (30%, n=10). This was followed by a decline at 60hr apf (26.7%, n = 30) and 72hr apf (11.8%, n = 85) reaching the lowest at eclosion (Fig S4A and S4B;1.7%, n=242). Thus, *mbl* expression in L4 neurons mirrors the expression of *Dscam2.10B*. Consistent with this, L2, L3 and L5, were all detected using the intersectional strategy with *Dscam2.10A-Gal4* but were not labelled using *mbl-Gal4* (Fig 3E). The expression of *mbl* is further strengthened by an independent RNA-seq study of isolated lamina neurons during development, where *mbl* is detected at high levels in L1, R7 and R8 neurons (~5-100 fold more than L2-L5)(Tan et al., 2015). Together, these results show *mbl* expression correlates with the cell-type-specific alternative splicing of *Dscam2.10B*. Importantly, this suggests that simply the presence or absence of *mbl* can determine the selection of the *Dscam2.10* isoform in a cell.

**Figure 3.**
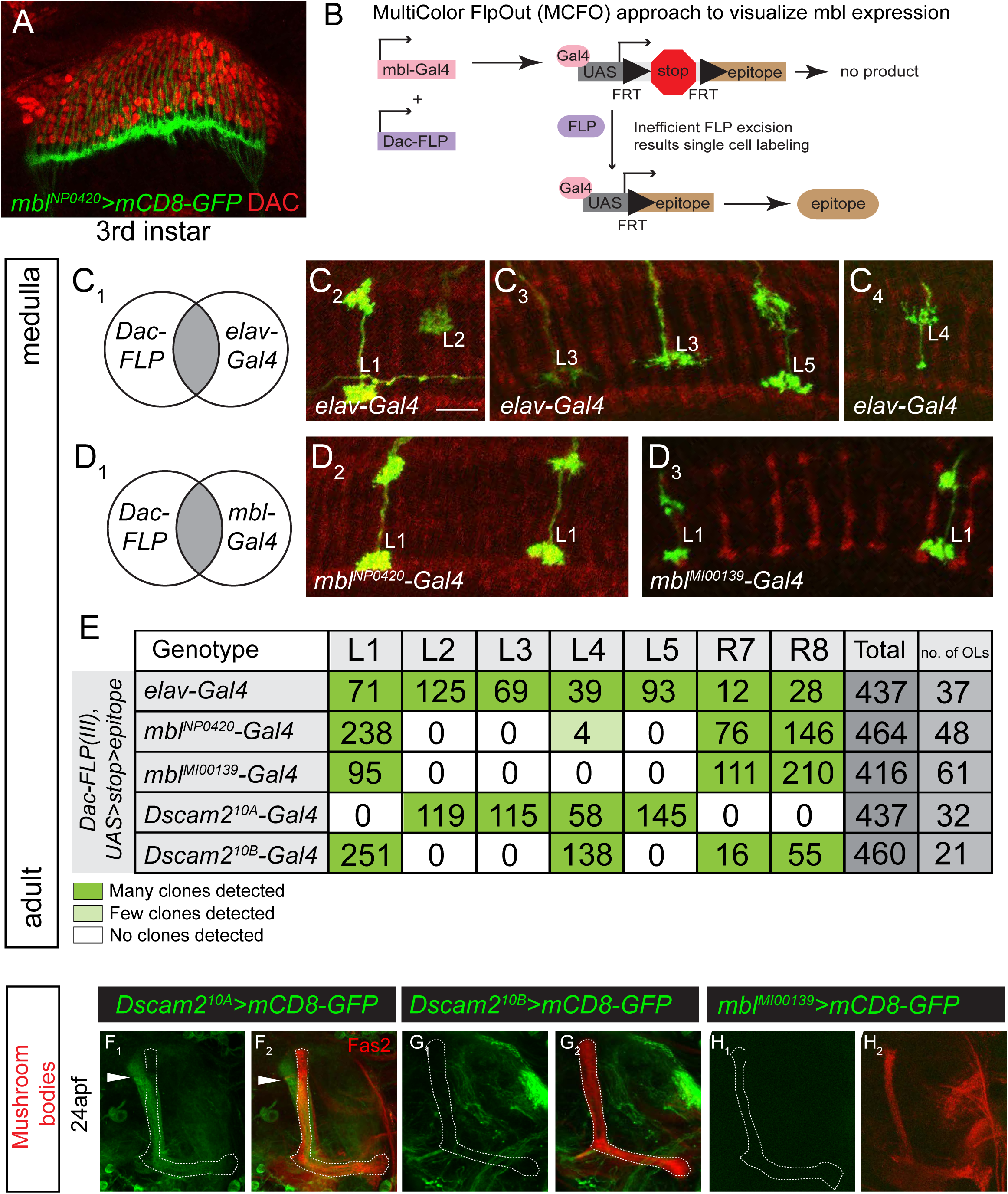
*mbl* is expressed in a cell-specific manner that correlates with *Dscam2.10B* (A) A *mbl*-*Gal4* reporter (green) is expressed in third instar R cells but not in lamina neuron precursor cells labelled with an antibody against Dacshund (DAC, red). (B) Schematic of MultiColor FlpOut (MCFO) approach to characterize *mbl* reporter expression in lamina neurons at adult stages. The UAS FlpOut construct produces an epitope-tagged version of a non-fluorescent GFP (smGFP,(Nern et al., 2015)) (C_1_-C_4_) All lamina neurons can be detected using a MCFO strategy with a pan-neuronal reporter (*elav-Gal4*). Lamina neurons were identified based on their unique axon morphologies. (D_1_-D_4_) An intersectional strategy using *mbl-Gal4* labels primarily L1 lamina neurons. (E) Quantification of lamina neurons and R7-R8 neurons observed using the intersectional strategy. Dark green and light green boxes represent high and low numbers of labelled neurons, respectively. (F-H) *Mbl* is not expressed in mushroom body (MB) neurons that express *Dscam2.10A* at 24hr apf. (F_1_-F_2_) *Dscam2.10A* is expressed in α’β’ MB neurons that are not labelled by Fas2. Fas2 labels the αβ and γ subsets of MB neurons. (G-H) Neither *Dscam2.10B* (G_1_-G_2_) nor *mbl* (H_1_-H_2_) are detected in MB neurons. See also Figures S3 and S4.

### Ectopic expression of multiple *mbl* isoforms is sufficient to promote the selection of *Dscam2* exon 10B

Since cells that select *Dscam2.10B* express *mbl* and cells that select *Dscam2.10A* lack *mbl*, we wondered whether it was sufficient to promote exon 10B selection in *Dscam2.10A*-positive cells. To test this, we ectopically expressed *mbl* with a ubiquitous driver (*Act5c-Gal4*) and monitored isoform B expression using *Dscam2.10B>tdTom*. We focussed on the mushroom body (MB), as this tissue expresses isoform A specifically in α’β’ neurons at 24hr apf where *mbl* is not detected (Fig 3G-3H, 4A-4C). Consistent with our prediction, ectopic expression of *mbl* using an enhancer trap containing a *UAS* insertion at the 5’ end of the gene (*Act5c>mbl*^*B2-E1*^), switched on *Dscam2.10B* in α’β’ MB neurons, where it is normally absent (Fig 4D). Driving *mbl* with a MB-specific *Gal4* (*OK107*) gave similar results (Fig 4E). Although our two *Gal4* drivers expressed *mbl* in all MB neurons, *Dscam2.10B* was only observed in α’β’ neurons, demonstrating that transcription of *Dscam2* is a pre-requisite for this splicing modulation. Previous studies have suggested that the *mbl* gene is capable of generating different isoforms with unique functions depending on their subcellular localization (Vicente et al., 2007). This also includes the production of a highly abundant circular RNA that can sequester the Mbl protein (Ashwal-Fluss et al., 2014; Houseley et al., 2006). To assess whether *Dscam2* exon 10B selection is dependent on a specific alternative variant of Mbl, we overexpressed the cDNAs of fly *mbl* isoforms (*mblA, mblB* and *mblC*; (Begemann et al., 1997; Juni and Yamamoto, 2009) as well as an isoform of the human *MBNL1* that lacks the linker region optimal for CUG repeat binding (*MBNL1_35_*; (Kino et al., 2004; Li et al., 2008) with either *Act5c-Gal4* or *OK107-Gal4*. These constructs all possess the tandem N-terminal CCCH motif that binds to YCGY sequences and lack the ability to produce *mbl* circRNA. In all cases, overexpression resulted in the misexpression of *Dscam2.10B* in α’β’ MBs (with the exception *Act5C>mblC*, which resulted in lethality; Fig 4D-4E). Using semi-quantitative RT-PCR from the *Act5C>mbl* flies, we demonstrated that overexpression of *mbl* did not lead to exon 10 skipping and that it increased exon 10B selection by 8-24% (Fig 4F), depending on the *mbl* isoform used. The inability of Mbl to completely inhibit exon 10A selection suggests that other factors or mechanisms may also contribute to cell-specific *Dscam2* isoform expression (see Discussion). These results suggest that Mbl protein isoforms are all capable of *Dscam2.10B* selection and independent of *mbl* circRNA. The ability of human MBNL1 to promote the selection of exon 10B suggests that the regulatory logic for *Dscam2* splicing is likely conserved in other mutually-exclusive cassettes in higher organisms. Together, our results show that all *mbl* isoforms are sufficient to promote *Dscam2.10B* selection.

**Figure 4.**
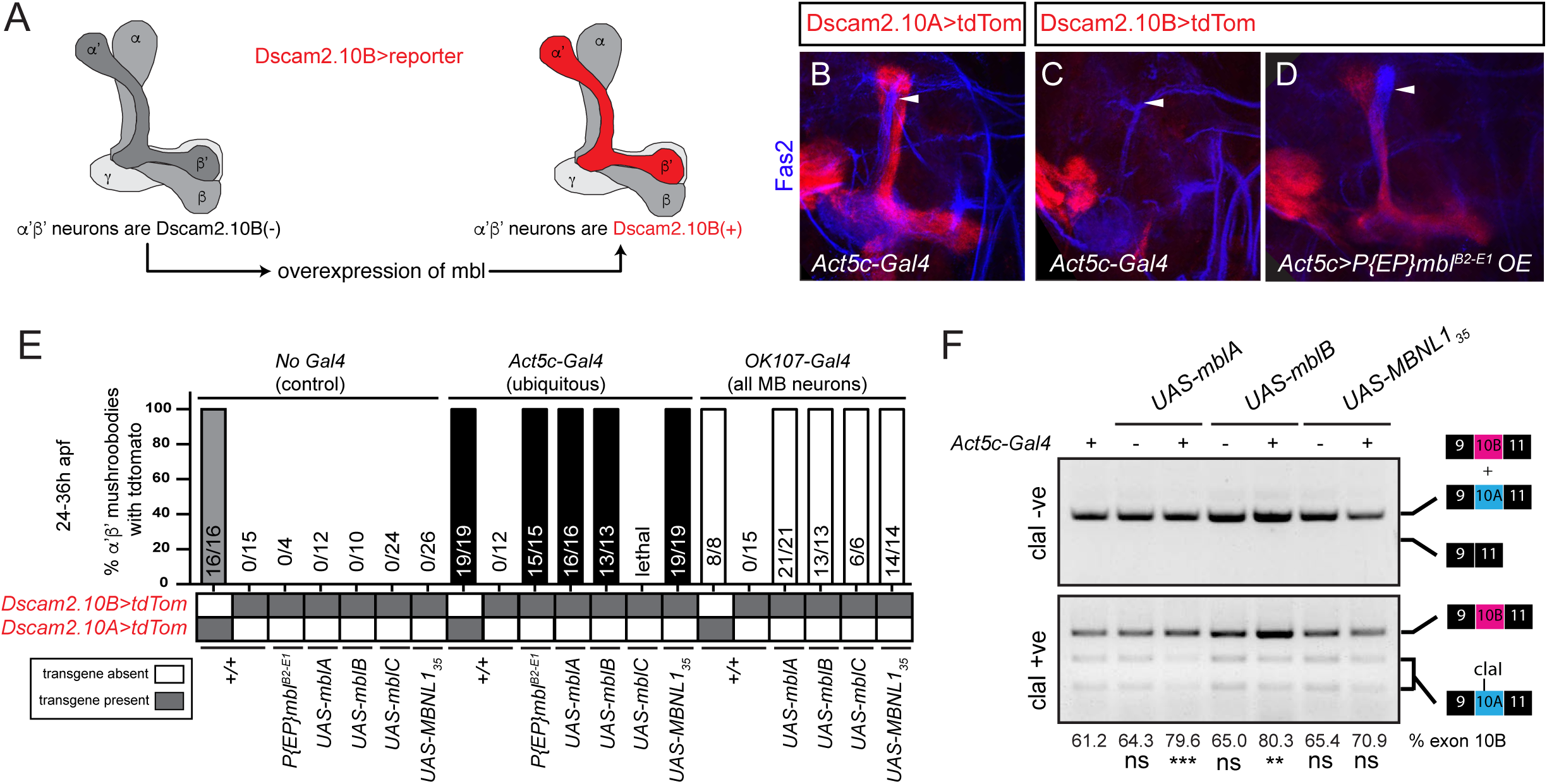
Multiple *mbl* isoforms promte selection of *Dscam2* exon 10B (A) Schematic showing that *mbl* is sufficient to drive *Dscam2.10B* selection in α’β’neurons. (B) Control showing that *Dscam2.10A* (red) is expressed in α’β’ neurons at 24hr apf. (C) Control showing that *Dscam2.10B* is normally repressed in α’β’ neurons. (D) Overexpression of *mbl* activates *Dscam2.10B* selection (red) in α’β’ neurons. (E) Quantification of *Dscam2.10* expression in α’β’ neurons at 24-36hr apf with and without *mbl* OE. Control (No Gal4, grey bar), ubiquitous driver (*Act5c-Gal4*, black bars) and pan-mushroom body neuron driver (*OK107-Gal4*, white bars). Y-axis represents the number of tdTom positive (+) α’β’ over the total, expressed as a percentage. Ratio of tdTom(+)/total is shown in each bar. (F) *Mbl* OE increases *Dscam2* exon 10B inclusion. Semiquantitative RT-PCR as in Figure 1. Exon 10A-specific cleavage with restriction enzyme ClaI shows an increase in exon 10B inclusion in *mbl* OE animals, without exon 10 skipping. Percentage of exon 10B inclusion was calculated by dividing 10B by 10A+10B bands following electrophoresis and densitometry. The mean of exon 10B inclusion is shown at the bottom of each lane. ANOVA test with Tukey’s multiple comparison test was used to compare the exon 10B inclusion. ns *P* > 0.05, ** *P* < 0.01, *** *P* < 0.001.

### Mbl regulates cell-type-specific *Dscam2* alternative splicing in lamina neurons

To determine whether the regulatory logic of *Dscam2* alternative splicing is consistent in other cell types, we manipulated *mbl* expression in lamina neurons (L1-L5). We first asked whether *mbl* LOF resulted in the de-repression of *Dscam2.10A* in L1 neurons. To do this, we visualized *Dscam2* isoform expression in L1-L5 using an intersectional strategy similar to Figure 3 but with a different *FLP* source (*27G05-FLP*). We detected L1 and L4 neurons when using the *Dscam2.10B-Gal4* reporter in a wild-type background, but not L2, L3 or L5. L1 was also not detected when using the *Dscam2.10A-Gal4* reporter, where L2-L5 cells were the primary neurons labelled (Fig 5A). Consistent with our R cell results, de-repression of *Dscam2.10A* was observed in L1 neurons in *mbl* transheterozygous animals (*mbl*^*e127*^*/mbl*^*MI00976*^) when compared to the corresponding heterozygous controls (*mbl*^*e127*^*/+* and *mbl*^*MI00976*^*/+,* Fig 5A-5B). We next asked whether ectopic overexpression of *mbl* would result in aberrant *Dscam.10B* selection in L2, L3 and L5 neurons where it is usually repressed. For this experiment, the *Gal4/UAS* system was used to overexpress *mbl* and the *LexA/LexAop* system was used to visualize *Dscam2* isoform expression. Using the same intersectional strategy, we found that *Dscam2-LexA* reporters showed similar patterns to the *Dscam2-Gal4* reporters (Fig 5C). Pan-neuronal overexpression (*elav-Gal4*) of *mbl* caused the aberrant detection of *Dscam2.10B* in L2, L3 and L5 cells that normally select *Dscam2.10A* (Fig 5C-5D). Together, our results show that Mbl regulates *Dscam2* cell-type-specific alternative splicing. Importantly, the simple presence or absence of *mbl* is sufficient to determine whether a cell expresses *Dscam2.10A* or *Dscam2.10B*.

**Figure 5.**
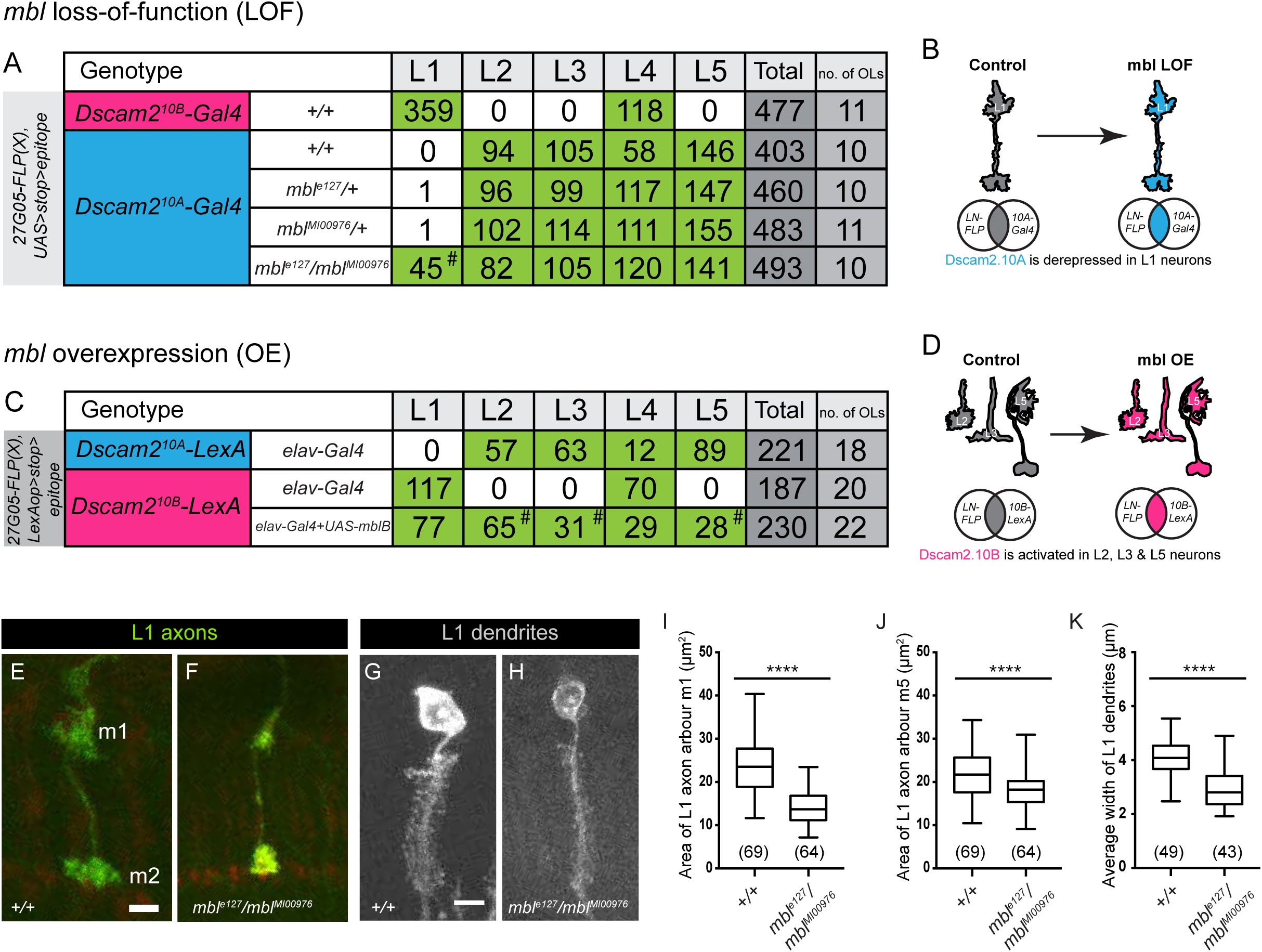
Mbl regulates *Dscam2* cell-type-specific alternative splicing in lamina neurons. (A) Quantification of lamina neurons L1-L5 observed using the *Dscam2.10B-Gal4* (magenta) or *Dscam2.10A-Gal4* (blue) reporters with the intersectional strategy in *mbl* LOF animals. Green boxes represent high number of labelled neurons. *Dscam2.10A* is de-repressed in L1 neurons in a *mbl* LOF background (*mbl*^*MI00976*^*/mbl*^*e27*^, hash tag). (B) Schematic of *Dscam2.10A* de-repression in *mbl* LOF L1 neurons. (C) Quantification of lamina neurons L1-L5 observed using the *Dscam2.10A-LexA* (blue) or *Dscam2.10B-LexA* (magenta) reporters with the intersectional strategy in animals with pan-neuronal (*elav-Gal4*) expression of *mbl*. Green boxes represent high numbers of labelled neurons. *Dscam2.10B-LexA* was aberrantly detected in L2, L3 and L5 neurons overexpressing *mblB* (hash tag). (D) Schematic of aberrant *Dscam2.10B* selection in L2, L3 and L5 neurons overexpressing *mbl*. (E-K) L1 neurons in *mbl* LOF animals have reduced axon arbor area and dendritic array width when compared to controls. (E) A representative confocal image of a control L1 axon (green) with arbors at m1 and m5 layers. (F) A representative confocal image of an L1 axon from *mbl* LOF animals (*mbl*^*MI00976*^*/mbl*^*e27*^). (G) A representative confocal image of a control L1 dendritic array (grey). (H) A representative confocal image of a L1 dendritic array from *mbl* LOF animals (*mbl*^*MI00976*^*/mbl*^*e27*^). (I) Quantification of L1 axon m1 arbor area (mm^2^). (J) Quantification of L1 axon m5 arbor area (mm^2^). (K) Quantification of L1 dendritic width (mm). Tukey boxplot format: middle line = median, range bars = min and max, box = 25–75% quartiles, and each data point = single cartridge. Numbers in parentheses represent total number of L1 neurons quantified. Parametric t-test was used to compare *mbl* LOF L1 axon arbour area with controls. Non-parametric t-test was used to compare *mbl* LOF L1 dendritic width with controls. **** *P*<0.0001. Bars, 5mm (E-H).

### Manipulation of *mbl* expression generates phenotypes observed in *Dscam2* single isoform mutants

If Mbl regulates *Dscam2* alternative splicing, *mbl* LOF and OE animals should exhibit similar phenotypes to *Dscam2* isoform misexpression. Previously, we showed that flies expressing a single isoform of *Dscam2* exhibit a reduction in L1 axon arbour size and well as reduced dendritic width (Kerwin et al., 2018; Lah et al., 2014). These flies were generated using recombinase-mediated cassette exchange and express a single isoform in all *Dscam2*-positive cells (Lah et al., 2014). The reduction in axonal arbors and dendritic widths were proposed to be due to inappropriate interactions between cells that normally express different isoforms. Consistent with these previous studies, we observed a reduction in the area of L1 axon arbors (more prominent in m1 than in m5, Fig 5E-5F and 5I-5J) and the width of dendritic arrays (Fig 5G-5H and 5G) in *mbl* transheterozygous animals (*mbl*^*e127*^*/mbl*^*MI00976*^) when compared to controls. Finally, we observed a phenotype in MB neurons overexpressing *mbl* where the β lobe neurons inappropriately crossed the midline (Fig S5A-S5C). Interestingly, a similar phenotype was observed in *Dscam2A* single isoform mutants. These data demonstrate that MB phenotypes generated in animals overexpressing *mbl*, phenocopy *Dscam2* single isoform mutants. While the origin of this non-autonomous phenotype is not known, it correlates with the misregulation of *Dscam2* alternative isoform expression.

## Discussion

In this study, we identify Mbl as a regulator of *Dscam2* alternative splicing. We demonstrate that removing *mbl* in a *mbl*-positive cell-type results in a switch from *Dscam2.10B* to *Dscam2.10A* selection. Ectopic expression of a variety of Mbl protein isoforms in a normally *mbl*-negative neuronal cell-type is sufficient to trigger the selection of *Dscam2.10B*. Consistent with this, transcriptional reporters demonstrate that *mbl* is expressed in a cell-type-specific manner in multiple cell-types, which tightly correlates with *Dscam2.10B*. Lastly, both *mbl* LOF and misexpression lead to phenotypes that are observed in flies that express a single *Dscam2* isoform.

Our data demonstrate that *mbl* is expressed in a cell-specific fashion. In the lamina of the fly visual system, L1 and L2 neurons are developmentally very similar in terms of both morphology and gene expression (Bausenwein et al., 1992; Fischbach and Dittrich, 1989; Tan et al., 2015). The difference in *mbl* expression between these two cells is critical for their development as when expression of this splicing factor is perturbed, both cells express the same isoforms and inappropriate Dscam2 interactions lead to phenotypes in their axons and dendrites. Although, cell-specific *mbl* expression has been alluded to previously (Huang et al., 2008; Machuca-Tzili et al., 2011; Norris et al., 2017), our study demonstrates that *mbl* regulation of *Dscam2* alternative splicing has functional consequences. Mbl appears to be regulated at the transcriptional level since the enhancer-trap as well as splicing-trap reporters lack the components crucial for post-transcriptional regulation yet still exhibit cell-type-specific expression (Fig 3). This was unexpected as a recent study showed that *mbl* encodes numerous alternative isoforms that could be individually post-transcriptionally repressed by different microRNAs, thus bypassing the need for transcriptional control of the gene. It will be interesting to explore the *in vivo* expression patterns of other splicing factors in *Drosophila* to determine whether cell-specific expression of a subset of splicing factors is a common mechanism for regulating alternative splicing in the brain.

The expression pattern of *mbl* and its ability to simultaneously repress exon 10A and select exon 10B suggest that this RNA binding protein and its associated co-factors are sufficient to regulate cell-type-specific splicing of *Dscam2*. *Dscam2.10A* could be the default exon selected when the Mbl complex is absent. In this way, cells that express *mbl* select *Dscam2.10B*. Consistent with this, ectopic expression of *mbl* in *mbl*-negative cells (L2, L3, L5 & α’β’ neurons) results in the aberrant selection of exon 10B. Our RT-PCR data, however, argue that *Dscam2* mutually exclusive alternative splicing may be more complicated than this model. Ubiquitous expression of *mbl* increased exon B inclusion modestly (up to 24%) as measured by RT-PCR (see Fig 4F). One might expect a more pronounced shift to isoform B if Mbl were the only regulator/mechanism involved. Further studies, including screens for repressors of exon 10B, will be required to resolve this issue.

The L1 axon and dendrite phenotypes generated through the LOF and ectopic expression of *mbl*, respectively, demonstrate that this splicing factor regulates aspects of neurodevelopment through cell-specific expression of *Dscam2* isoforms. In the lamina, *mbl* expression in L1, and its absence in L2, permits these neurons to express distinct Dscam2 proteins that cannot recognise each other. Phenotypes arise in these neurons both when they are engineered to express the same isoform (Kerwin et al., 2018; Lah et al., 2014) and when mbl is misregulated (Fig 5). These data strongly link the regulation of cell-specific *Dscam2* splicing with normal neuron development. *Mbl* overexpression also generates a midline crossing phenotype in MB neurons that is similar to that observed in animals expressing a single isoform. This phenotype is complicated, however, by the observation that *Dscam2.10A*, but not *Dscam2.10B*, animals show a statistically significant increase in midline crossing compared to controls (Fig S4). This issue may have to do with innate differences between isoform A and isoform B that are not completely understood. It is possible that isoform A and B are not identical in terms of signalling due to either differences in homophilic binding or differences in co-factors associated with specific isoforms. Consistent with this notion, we previously reported that *Dscam2.10A* single isoform lines produce stronger phenotypes at photoreceptor synapses compared to *Dscam2.10B* (Kerwin et al., 2018).

Together, our results demonstrate that the simple presence or absence of a splicing factor can affect neurodevelopment through the cell-specific selection of distinct isoforms of a cell surface protein. Although we provide compelling genetic evidence of how Mbl regulates the alternative splicing of *Dscam2*, the regulatory logic we discovered for *Dscam2* is likely to extend to cover the splicing events of many other genes crucial for neurodevelopment. Developmental analysis of *mbl* expression in the cells studied here suggests that it turns on after neurons have obtained their identity (similar to *Dscam2*) and is therefore well suited for regulating processes such as axon guidance and synapse specification. Identifying these splicing events may provide clues into how the brain can diversify and regulate its repertoire of proteins to promote neural connectivity.

## Experimental procedures

### Fly strains

*Dscam2.10A-LexA* and *Dscam2.10B-LexA* (Tadros et al., 2016), *UAS-Dcr2* and *UAS-mbl-RNAi*^*VDRC28732*^(Dietzl et al., 2007), *LexAop-myr-tdTomato* (attP2, (Chen et al., 2014), *UAS-Srp54-RNAi*^*TRiP.HMS03941*^, *CadN-RNAi*^*TRiP.HMS02380*^and *UAS-mbl-RNAi*^*TRiP.JF03264*^(Ni et al., 2008), *UAS-mCD8-GFP* (Lee and Luo, 1999), *FRT42D* (Xu and Rubin, 1993), *mbl*^*e127*^and *mbl*^*e27*^(Begemann et al., 1997), *mbl*^*MI00976*^and *mbl*^*MI04093*^(Venken et al., 2011), *Df(2R)BSC154* (Cook et al., 2012), *Df(2R)Exel6066* (Parks et al., 2004), *ey-FLP* (Chr.1, (Newsome et al., 2000), *GMR-myr-GFP, mbl*^*NP0420*^*-Gal4* and *mbl*^*NP1161*^*-Gal4* (Hayashi et al., 2002), *mbl*^*k01212*^*-LacZ* (Spradling et al., 1999), *mbl*^*MiMIC00139*^*-Gal4* (H. Bellen Lab), *Dac-FLP* (Chr.3, (Millard et al., 2007),*UAS>stop>myr::smGdP-V5-THS-UAS>stop>myr::smGdP-cMyc* (attP5, (Nern et al., 2015), *Dscam2.10A-Gal4* and *Dscam2.10B-Gal4* (Lah et al., 2014) *Act5C-Gal4* (Chr.3, from Yash Hiromi), *OK107-Gal4* (Connolly et al., 1996), *UAS-mblA, UAS-mblB* and *UAS-mblC* (D. Yamamoto Lab), *P{EP}mbl*^*B2-E1*^, *UAS-mblA-FLAG* and *UAS-MBNL1*_*35*_(Li et al., 2008).

### RNAi screening

The RNAi screen line was generated as follows: *GMR-Gal4* was recombined with *GMR-GFP* on the second chromosome. *Dscam2.10A-LexA* (Tadros et al. 2016) was recombined with *LexAop-myr-tdTomato* on the third chromosome. These flies were crossed together with *UAS-Dcr-2* (X) to make a stable RNAi screen stock. UAS-RNAi lines were obtained from Bloomington and VDRC. Lethal UAS-RNAi stocks were placed over balancers with developmentally selectable markers. Virgin females were collected from the RNAi screen stock, crossed to UAS-RNAi males and reared at 25°C. Wandering third instar larvae were dissected and fixed. We tested between one to three independent RNAi lines per gene. In total, we imaged ~2300 third instar optic lobes without antibodies using confocal microscopy at 63X. RNAi lines tested are listed in Table S1.

### Semiquantitative and quantitative RT-PCR

Total RNA was isolated using TRIzol (Ambion) following the manufacturer’s protocol. Reverse transcription was performed on each RNA sample with random primer mix (semiquantitative, NEB) or Oligo-dT (qRT-PCR, NEB) using 200 units of M-MULV (NEB) and 1 μg of RNA in a 20 μL reaction, at 42°C for 1 hr. PCR reactions were set up with specific primers to analyse alternative splicing of various regions of *Dscam2*. Where possible, semi-quantitative PCR was performed to generate multiple isoforms in a single reaction and relative levels were compared by electrophoresis followed by densitometry. For qRT-PCR, 1μL of CDNA were added to a Luna Universal SYBR-Green qPCR Master Mix kit (NEB). Samples were added into a 200μL 96-well plate and read on the QuantStudio TM 6 Flex Real-Time PCR machine. Rq values were calculated in Excel (Microsoft).

### Immunohistochemistry

Immunostaining were conducted as previously described (Lah et al. 2014). Antibody dilutions used were as follows: mouse mAb24B10 (1:20; DSHB), mouse anti-Repo (1:20; DSHB), mouse anti-DAC (1:20; DSHB), mouse anti-Fas2 (1:20; DSHB) rat anti-ELAV (1:200), V5-tag:DyLight anti-mouse 550 (1:500; AbD Serotec), V5-tag:DyLight anti-mouse 405 (1:200; AbD Serotec), myc-tag;DyLight anti-mouse 549 (1:200; AbD Serotec), Phalloidin:Alexa Fluor 568 (1:200; Molecular Probes), DyLight anti-mouse 647 (1:2000; Jackson Laboratory), DyLight Cy3 anti-rat (1:2000; Jackson Laboratory).

### Image acquisition

Imaging was performed at the School of Biomedical Sciences Imaging Facility. Images were taken on a Leica SP8 laser scanning confocal system with a 63X Glycerol NA 1.3.

### Fly genotypes

Specific genotypes can be found in the supplemental text.

### Author contribution

J.S.S.L designed and performed all experiments. S.S.M supervised the project. J.S.S.L and S.S.M wrote the manuscript.

## Acknowledgements

We thank Wael Tadros, Yi Chen, Larry Zipursky, Greg Neely, Louis O’Keefe, Nancy Bonini, Aljoscha Nern and Bloomington Stock Center for sharing fly stocks. We thank the Daisuke Yamamoto Lab for constructing the *UAS-mbl* lines deposited and maintained at the Kyoto Stock Center. We thank Shaun Walters for technical assistance on the Leica confocal microscopy. We note that Grace Shin initially observed *Dscam2* isoform expression in the adult mushroom bodies. We thank Kevin Mutemi for his thorough characterization of *Dscam2* isoform expression in mushroom bodies during development and all midline crossing defects in *Dscam2* single isoform mutant animals. We thank Wei Jun Tan for the heroic feat of triple balancing *OK107-Gal4*. We also thank members of the Millard, Pecot, Hilliard and van Swinderen lab for their feedback. The RNAi screen was inspired by the works of Hidehito Kuroyanagi. This work was supported by the National Health and Medical Research Council of Australia (NHMRC grant APP1021006). J.S.S.L was supported by the Australia Postgraduate Award (Research Training Scheme) from the Australian Federal Government and the Lavidis grant in aid.

## Fly genotypes

### R cell RNAi experiments (Figure 1)

1. *w; GMR-GFP, GMR-Gal4/CyO; Dscam2.10B-LexA, LexAop-myr-tdTomato/TM6B*
2. *w; GMR-GFP, GMR-Gal4/CyO; Dscam2.10A-LexA, LexAop-myr-tdTomato/TM6B*
3. *w, UAS-Dcr-2; GMR-GFP, GMR-Gal4/CyO; Dscam2.10A-LexA, LexAop-myr-tdTomato/TM6B*
4. *w, UAS-Dcr-2; GMR-GFP, GMR-Gal4/UAS-mCD8-RFP; Dscam2.10A-LexA, LexAop-myr-tdTomato/+*
5. *w, UAS-Dcr-2; GMR-GFP, GMR-Gal4/UAS-mbl-RNAi(v28732); Dscam2.10A-LexA, LexAop-myr-tdTomato/+*
6. *w, UAS-Dcr-2; GMR-GFP, GMR-Gal4/+; Dscam2.10A-LexA, LexAop-myr-tdTomato/UAS-mbl-RNAi(TRiP.JF03264)*

### mbl whole animal experiments (Figure 1)

1. *w; +; Dscam2.10B-LexA, LexAop-myr-tdTomato/TM6B*
2. *w; +; Dscam2.10A-LexA, LexAop-myr-tdTomato/TM6B*
3. *w; mbl*^*e127*^*/CyO,GFP; Dscam2.10A-LexA, LexAop-myr-tdTomato/TM6B*
4. *w; mbl*^*MI00976*^*/CyO,GFP; Dscam2.10A-LexA, LexAop-myr-tdTomato/TM6B*
5. *w; mbl*^*MI04093*^*/CyO,GFP; Dscam2.10A-LexA, LexAop-myr-tdTomato/TM6B*
6. *w; mbl*^*e127*^*/ mbl*^*MI00976*^; *Dscam2.10A-LexA, LexAop-myr-tdTomato/+*
7. *w; mbl*^*MI04093*^*/ mbl*^*MI00976*^; *Dscam2.10A-LexA, LexAop-myr-tdTomato/+*

### mbl ey-FLP MARCM experiments (Figure 2)

1. *w, ey-FLP; FRT42D, Tub-Gal80/FRT42D; Dscam2.10A-LexA, LexAop-myr-tdTomato, Act5c-Gal4, UAS-mCD8-GFP/+*
2. *w, ey-FLP; FRT42D, Tub-Gal80/FRT42D, mbl*^*e27*^; *Dscam2.10A-LexA, LexAop-myr-tdTomato, Act5c-Gal4, UAS-mCD8-GFP/+*
3. *w, ey-FLP; FRT42D, Tub-Gal80/FRT42D, mbl*^*e127*^; *Dscam2.10A-LexA, LexAop-myr-tdTomato, Act5c-Gal4, UAS-mCD8-GFP/+*

### mbl expression experiments (Figure 3)

1. *w; UAS-mCD8-GFP/+; mbl*^*NP0420*^*-Gal4/+*
2. *w; UAS-mCD8-GFP/+; mbl*^*MI00139*^*-Gal4/+*
3. *w; Dac-FLP/+; elav-Gal4/ UAS>stop>myr::smGdP-V5-THS-UAS>stop>myr::smGdP-cMyc*
4. *w; Dac-FLP/+; mbl*^*NP0420*^*-Gal4/ UAS>stop>myr::smGdP-V5-THS-UAS>stop>myr::smGdP-cMyc*
5. *w; Dac-FLP/+; mbl*^*MI00139*^*-Gal4/ UAS>stop>myr::smGdP-V5-THS-UAS>stop>myr::smGdP-cMyc*
6. *w; Dac-FLP/+; Dscam2.10A-Gal4/ UAS>stop>myr::smGdP-V5-THS-UAS>stop>myr::smGdP-cMyc*
7. *w; Dac-FLP/+; Dscam2.10B-Gal4/ UAS>stop>myr::smGdP-V5-THS-UAS>stop>myr::smGdP-cMyc*
8. *w; +; mbl*^*NP0420*^*-Gal4/UAS-GFP.nls*
9. *w; +; mbl*^*MI00139*^*-Gal4/UAS-GFP.nls*

### mbl ectopic expression in MBs (Figure 4)

1. *w; +; Dscam2.10A-LexA, LexAop-myr-tdTomato, Act5c-Gal4, UAS-mCD8-GFP/+*
2. *w; +; Dscam2.10B-LexA, LexAop-myr-tdTomato, Act5c-Gal4, UAS-mCD8-GFP/+*
3. *w; P{EP}mbl*^*B2-E1*^*/+; Dscam2.10B-LexA, LexAop-myr-tdTomato, Act5c-Gal4, UAS-mCD8-GFP/+*
4. *w; +; Dscam2.10B-LexA, LexAop-myr-tdTomato, Act5c-Gal4, UAS-mCD8- GFP/UAS-mblA*
5. *w; +; Dscam2.10B-LexA, LexAop-myr-tdTomato, Act5c-Gal4, UAS-mCD8- GFP/UAS-mblB*
6. *w; +; Dscam2.10B-LexA, LexAop-myr-tdTomato, Act5c-Gal4, UAS-mCD8- GFP/UAS-mblC*
7. *w; +; Dscam2.10B-LexA, LexAop-myr-tdTomato, Act5c-Gal4,UAS-mCD8- GFP/UAS-MBNL1_35_*
8. *w; +; Dscam2.10B-LexA, LexAop-myr-tdTomato, UAS-mCD8-GFP/UAS-mblA; OK107-Gal4/+*
9. *w; +; Dscam2.10B-LexA, LexAop-myr-tdTomato, UAS-mCD8-GFP/UAS-mblB; OK107-Gal4/+*
10. *w; +; Dscam2.10B-LexA, LexAop-myr-tdTomato, UAS-mCD8-GFP/UAS-mblC; OK107-Gal4/+*
11. *w; +; Dscam2.10B-LexA, LexAop-myr-tdTomato, UAS-mCD8-GFP/UAS- MBNL1_35_; OK107-Gal4/+*

### Lamin neuron FlpOut mbl LOF (Figure 5)

1. *w, 27G05-FLP/(+ or Y); Bl/CyO; Dscam2.10B-Gal4/ UAS>stop>myr::smGdP-V5- THS-UAS>stop>myr::smGdP-cMyc*
2. *w, 27G05-FLP/(+ or Y); Bl/CyO; Dscam2.10A-Gal4/ UAS>stop>myr::smGdP-V5- THS-UAS>stop>myr::smGdP-cMyc*
3. *w, 27G05-FLP/(+ or Y); mbl*^*e127*^*/ CyO; Dscam2.10A-Gal4/ UAS>stop>myr::smGdP-V5-THS-UAS>stop>myr::smGdP-cMyc*
4. *w, 27G05-FLP/(+ or Y); mbl*^*MI00976*^*/CyO; Dscam2.10A-Gal4/ UAS>stop>myr::smGdP-V5-THS-UAS>stop>myr::smGdP-cMyc*
5. *w, 27G05-FLP/(+ or Y); mbl*^*e127*^*/ mbl*^*MI00976*^; *Dscam2.10A-Gal4/ UAS>stop>myr::smGdP-V5-THS-UAS>stop>myr::smGdP-cMyc.*
6. *w, 27G05-FLP/(+ or Y); elav-Gal4/LexAop2>stop>myr::smGdP-V5; Dscam2.10A-LexA/TM2.*
7. *w, 27G05-FLP/(+ or Y); elav-Gal4/LexAop2>stop>myr::smGdP-V5; Dscam2.10B-LexA/TM2.*
8. *w, 27G05-FLP/(+ or Y); elav-Gal4/LexAop2>stop>myr::smGdP-V5; Dscam2.10B-LexA/UAS-mblB.*

### L1 axonal and dendritic defects (Figure 5)

1. *w, 27G05-FLP/(+ or Y); Bl.CyO; Dscam2.10A-Gal4/ UAS>stop>myr::smGdP-V5- THS-UAS>stop>myr::smGdP-cMyc.*
2. *w, 27G05-FLP/(+ or Y); mbl*^*e127*^*/ mbl*^*MI00976*^; *Dscam2.10A-Gal4/ UAS>stop>myr::smGdP-V5-THS-UAS>stop>myr::smGdP-cMyc.*

### mbl ey-FLP mosaic experiments (Figure S1)

1. *w, ey-FLP; FRT42D, GMR-myr-GFP/FRT42D; Dscam2.10B-LexA, LexAop-myr-tdTomato, UAS-mCD8-GFP/+*
2. *w, ey-FLP; FRT42D, GMR-myr-GFP/FRT42D; Dscam2.10A-LexA, LexAop-myr-tdTomato, UAS-mCD8-GFP/+*
3. *w, ey-FLP; FRT42D, GMR-myr-GFP/FRT42D, Df(2R)154; Dscam2.10A-LexA, LexAop-myr-tdTomato, UAS-mCD8-GFP/+*
4. *w, ey-FLP; FRT42D, GMR-myr-GFP/FRT42D, mbl*^*e27*^; *Dscam2.10A-LexA, LexAop-myr-tdTomato, UAS-mCD8-GFP/+*
5. *w, ey-FLP; FRT42D, GMR-myr-GFP/FRT42D, mbl*^*MI00976*^; *Dscam2.10A-LexA, LexAop-myr-tdTomato, UAS-mCD8-GFP/+*

### mbl expression (Figure S3)

1. *w; mbl*^*K01212*^*-LacZ*
2. *w; mbl*^*NP1161*^*-Gal4/CyO, UAS-LacZ*
3. *w; mbl*^*MI00139*^*-Gal4/+; UAS-CD8-GFP/+*
4. *w; mbl*^*MI00139*^*-Gal4/UAS-GFP.nls*
5. *w; mbl*^*NP0420*^*-Gal4/UAS-GFP.nls*

### MB axon defects (Figure S5)

1. *w; +; +*
2. *w; +; Dscam2*^*null*^*/ Dscam2*^*null*^
3. *w; +; Dscam2A/ Dscam2A*
4. *w; +; Dscam2B/ Dscam2B*
5. *w; mbl*^*e127*^*/ mbl* ^*MI0097*^
6. *w; +; +; OK107-Gal4/+*
7. *w; UAS-mbl-RNAi(v28732)/+*
8. *w; UAS-mbl-RNAi(v28732)/+; +; OK107-Gal4/+*
9. *w; P{EP}mbl*^*B2-E1*^*/+*
10. *w; P{EP}mbl*^*B2-E1*^*/+; +; OK107-Gal4/+*
11. *w; +; UAS-mblA/+*
12. *w; +; UAS-mblA/+; OK107-Gal4/+*
13. *w; +; UAS-mblB/+*
14. *w; +; UAS-mblB/+; OK107-Gal4/+*
15. *w; +; UAS-mblC/+*
16. *w; +; UAS-mblC/+; OK107-Gal4/+*
17. *w; +; UAS-MBNL1_35_/+*
18. *w; +; UAS-MBNL1_35_/+; OK107-Gal4/+*

**Figure S1.**
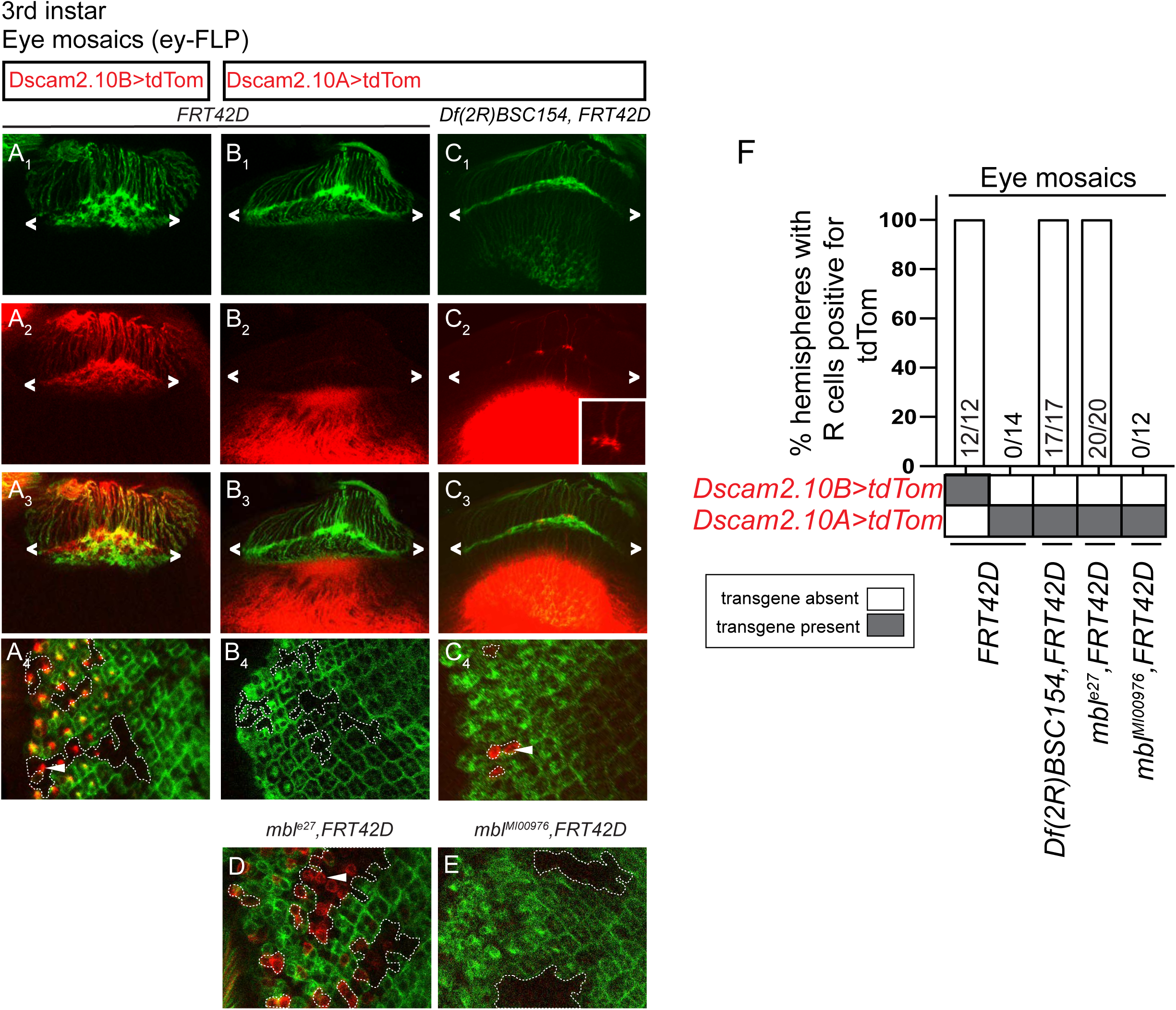
Related to Figure 1. *Mbl* LOF results in aberrant *Dscam2.*10A reporter expression in eye mosaic clones. (A-F) Eye mosaics of *mbl* LOF alleles cause de-repression of *Dscam2.10A>tdTom* in R cells. *WT* mosaic clones (GFP-negative) express *Dscam2.10B>tdTom* (A_1_-A_4_) but not *Dscam2.10A>tdTom* (B_1_-B_4_). *Mbl* mutant (GFP-negative) clones, *Df(2R)BSC154* show aberrant *Dscam2.10A* expression in R cells (C -C). (D) mbl^e27^eye clones exhibit de-repression of Dscam2.10A (red). (E) Clones of a *mbl* allele that deleted only a portion of all mbl isoforms (*mbl*^*MI00976*^) do not exhibit de-repression of Dscam2.10A. (F) Quantification of *Dscam2.10>tdTom* expression in third instar R cells with *mbl* LOF eye mosaic clones. Y-axis represents the number of optic lobes with R cells positive for tdTom over total number of optic lobes quantified as a percentage. On the x-axis, the presence of a transgene is indicated with a grey box.

**Figure S2.**
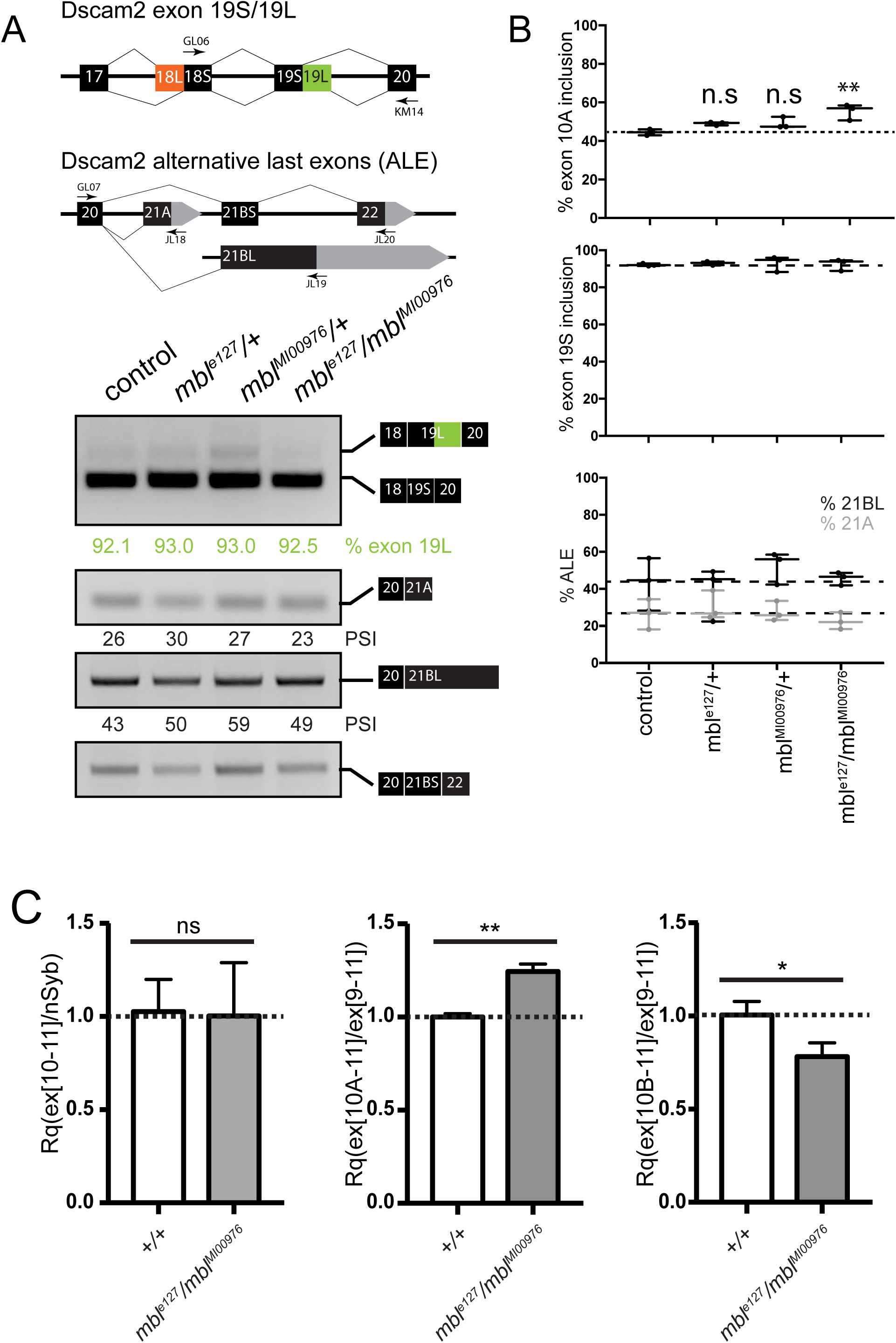
Related to Figure 1. *Mbl* LOF is associated with increased *Dscam2.10A* inclusion without affecting other *Dscam2* splicing events. (A) *Mbl* LOF (*mbl*^*e127*^/*mbl*^*MI00976*^) does not affect other *Dscam2* splicing events. Semiquantitative RT-PCR from different genotypes indicated. Primers amplified the variable region that includes exon 19S/19L or three alternative last exons (ALE). Percentage of 19L inclusion was calculated by dividing the 19L band by 19L+19S. Percentage of ALE 21A and ALE 21BL inclusion was calculated by dividing respectively the 21A and 21BL band by 21A+21BL+21BS (total). (B) Graphs of RT-PCR data from A and Figure 1P. Top graph depicts Dscam2.10A inclusion. Middle graph represents exon 19S inclusion. Bottom graph represents percentage inclusion of different ALEs. Plots show minimum (bottom line), mean (middle line) and maximum (top line) points, where individual points depict biological replicates. Dashed line represents mean of control. (C) Quantitative RT-PCR of *mbl* LOF mutant (*mbl*^*e127*^/*mbl*^*MI00976*^) show increased exon 10A inclusion and decreased exon 10B inclusion. The left graph shows *Dscam2.10* levels compared to *synaptobrevin* (*nSyb*). The middle graph shows *Dscam2.10A* levels compared to *Dscam2.10*. The right graph shows *Dscam2.10B* levels compared to *Dscam2.* Bar graph format (error bars depict standard error of means). The y-axis is the relative quantity (Rq). Dashed line represents mean of control. Unpaired t-test was conducted to compare Rq levels between control and *mbl* LOF mutants. ns *P* > 0.05, * *P* < 0.05, ** *P* < 0.01.

**Figure S3.**
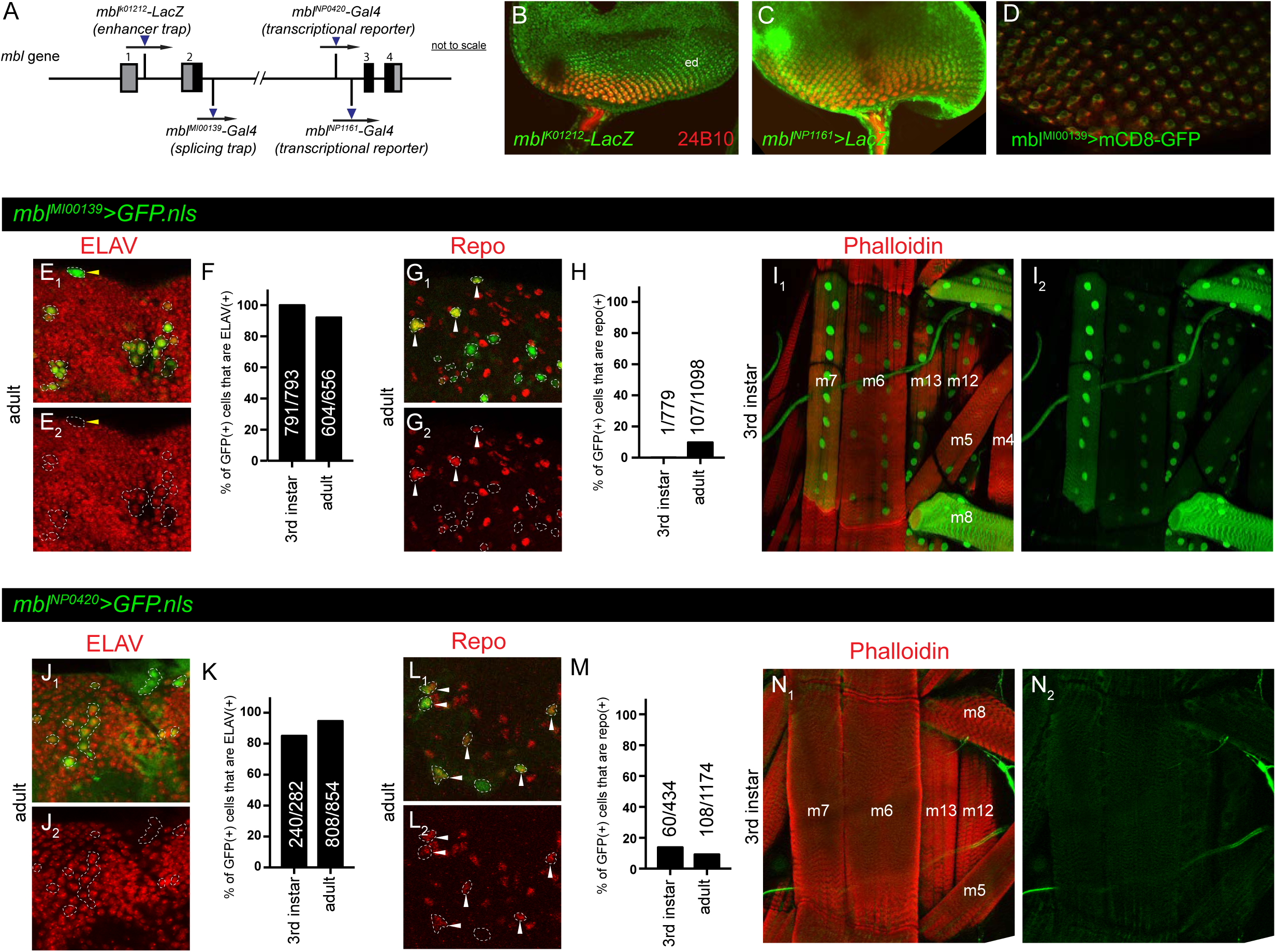
Related to Figure 3. *Mbl* is expressed in R cells, neurons and glia (A) Schematic showing the insertion locations of different *mbl* reporters. Translated regions (black) and non-translated regions (grey) are shown. (B-D) *Mbl* is expressed in R cells (red) in third instar eye-discs (ed). The *mbl* enhancer traps *mbl*^*K01212*^*-LacZ* (B), *mbl*^*NP1161*^*-Gal4* (C) and splicing trap reporter *mbl*^*MI00139*^*-Gal4* (D, green) overlapped with a marker of R cells (24B10). (E-I) *mbl*^*MI00139*^*>GFP.nls* is expressed in neurons and muscles. (E_1_ –E_2_) Representative confocal image of a *mbl*^*MI00139*^*>GFP.nls* (green) adult brain co-labelled with an ELAV antibody (red). Dashed lines demarcate GFP(+) cells. Yellow solid arrowheads show GFP(+) cells that are ELAV(-). (F) Quantification of *mbl* in third instar and adult brains where ~90-100% of GFP(+) cells are also ELAV(+) (black bars). Y-axis represents the number of GFP(+) cells positive for ELAV quantified as a percentage. (G_1_ -G_2_) Representative confocal image of a *mbl*^*MI00139*^*>GFP.nls* adult brain labelled with a Repo antibody (red). Dashed lines demarcate GFP(+) cells. White solid arrowheads show GFP(+) cells that are positive for Repo. (H) Quantification of *mbl*^*MI00139*^*>GFP.nls* where ~0-10% of *mbl*^*MI00139*^*>GFP.nls* (+) cells are also Repo(+).Y-axis represents the number of GFP(+) cells positive for Repo quantified as a percentage. (I -I) *mbl*^*MI00139*^*>GFP.nls* expression is also found in third instar muscles m4-m8, m12 and m13 (Phalloidin, red). (J_1_ -J_2_) Representative confocal image of a *mbl*^*NP0420*^*>GFP.nls* (green) adult brain co-labelled with an ELAV antibody (red). Dashed lines demarcate GFP(+) cells. (K) Quantification of *mbl*^*NP0420*^*>GFP.nls* in third instar and adult brains where ~80-90% of GFP(+) cells are also ELAV(+). (L-M) In third instar and adult brains, *mbl*^*NP0420*^*>GFP.nls* overlaps minimally with Repo (red). (L -L) Representative confocal image of a *mbl*^*NP0420*^*>GFP.nls* adult brain labelled with Repo. Dashed lines demarcate GFP(+) cells. White solid arrowheads show GFP(+) cells that are positive for Repo. (M) Quantification of *mbl*^*NP0420*^*>GFP.nls* in third instar and adult brains where ~10-15% of GFP (+) cells are also Repo(+). (N -N) *mbl*^*NP0420*^*>GFP.nls* expression is not detected in third instar muscles m4-m8, m12 and m13 (Phalloidin, red).

**Figure S4.**
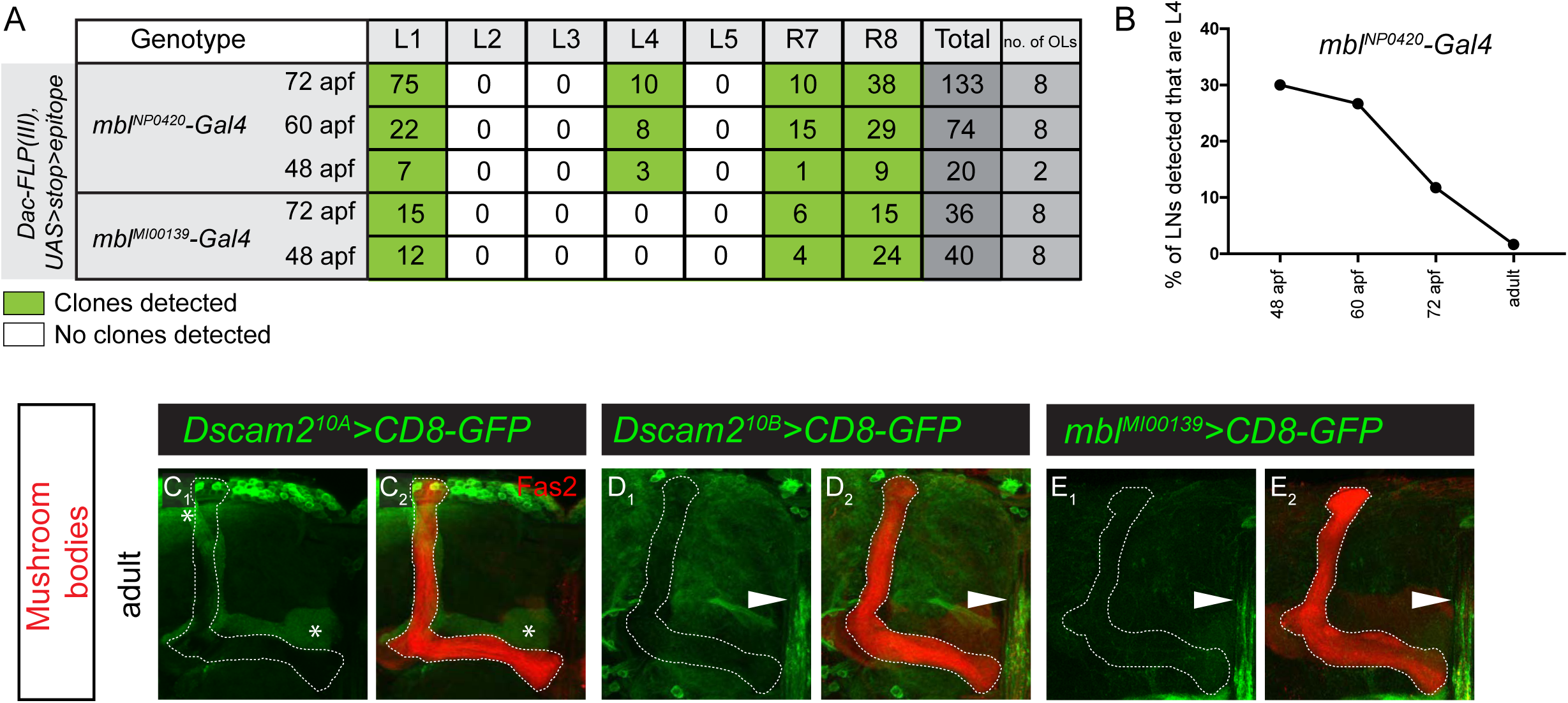
Related to Figure 3. *Mbl* expression is cell-type-specific and correlates with *Dscam2.10B*. (A) Quantification of lamina neurons and R7-R8 neurons observed using the intersectional strategy during development. Two different *mbl* reporters were used. The transcriptional reporter labelled L4 cells early in development whereas the splicing trap reporter did not. This is most likely due to the lower efficiency of the splicing trap given that it produced 5X fewer L1 clones at 72hr compared to the transcriptional reporter. Green boxes represent detection of reporter expression at different hours after pupal formation (apf). (B) A plot of the percentage of L4 lamina neurons over total lamina neurons during development (data from the *mbl* transcriptional reporter). (C-E) *Mbl* is not detected in MB neurons that express *Dscam2.10A* in adults. (C_1_-C_2_) *Dscam2.10A* is expressed in α’β’ mushroom body neurons (asterisks) but not the αβ and g subsets of MB neurons labelled by Fas2 (red). Neither *Dscam2.10B* (D_1_-D_2_) nor *mbl* (E_1_-E_2_) are expressed in MB neurons. Neurons in the midline express both *Dscam2.10B* and *mbl* (white arrowhead).

**Figure S5.**
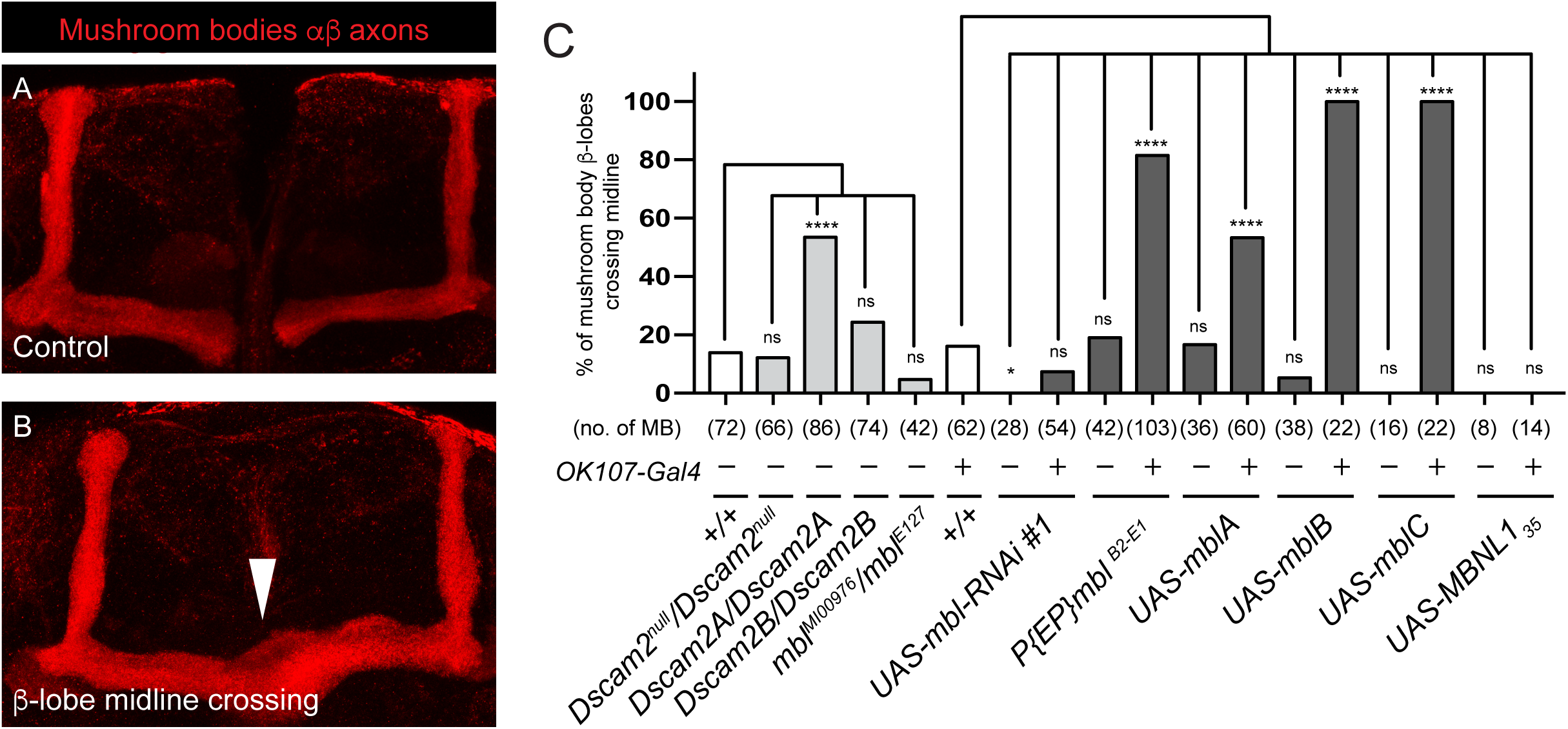
Related to Figure 4. Neurons overexpressing *mbl* phenocopy *Dscam2* single isoform mutants (A-B) MBs overexpressing *mbl* exhibit defects associated with *Dscam2* single isoform mutants. (A) A representative confocal image of control adult αβ lobes (red) with clear separation between the two β-lobes at the midline. (B) A representative confocal image of adult αβ lobes from an animal overexpressing *mblA*. b-lobe axons inappropriately cross the midline (arrowhead). (C) Quantification of β-lobe axon midline crossing defects. Numbers in parentheses represent total number of MBs quantified. Fishers exact test was used to compare genotypes to their corresponding controls (white bars). ns (not significant) *P*>0.05, * *P*<0.05 and **** *P*<0.0001.

**Table S1.**
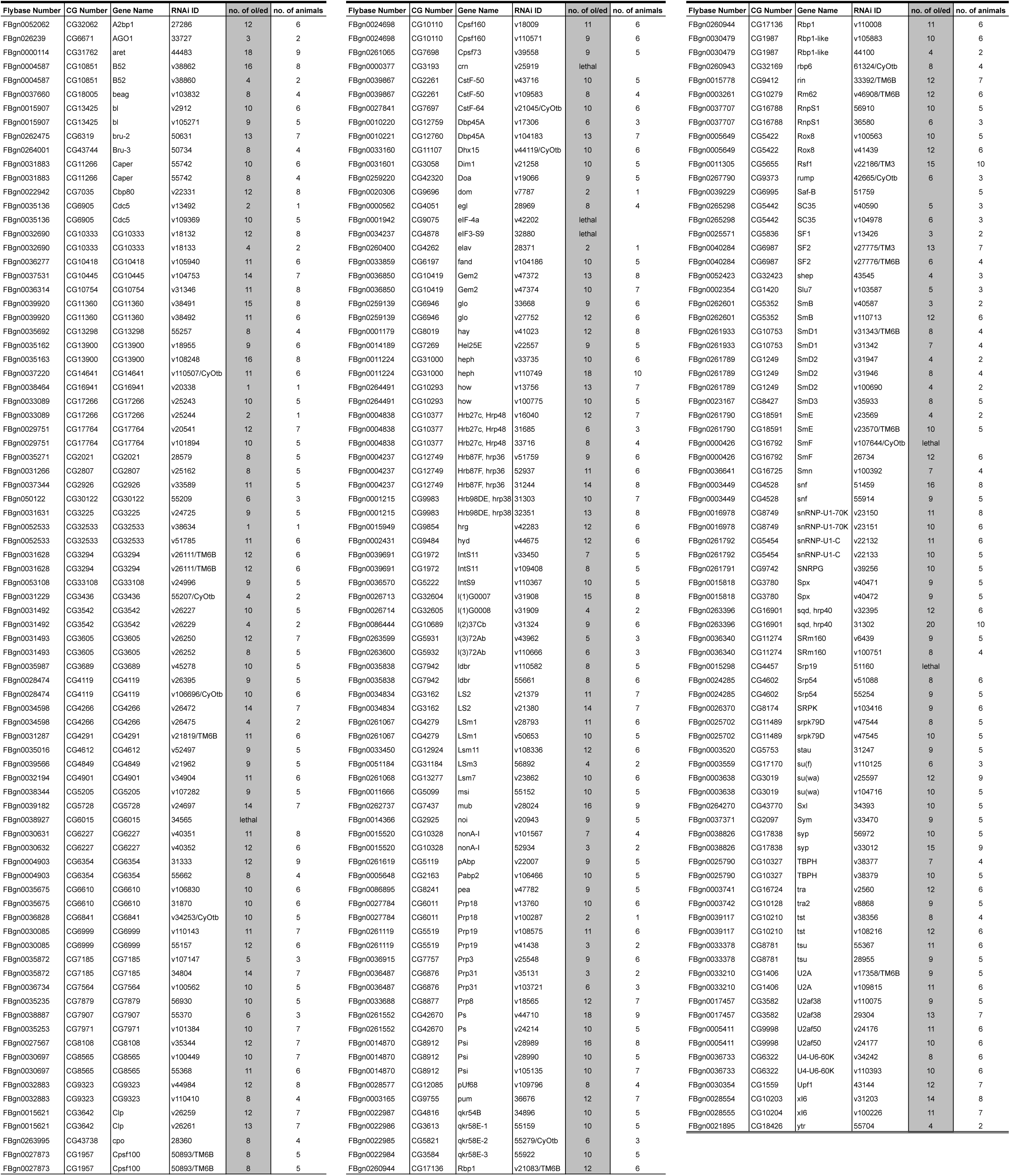
Related to Figure 1. List of tested RNAi that did not de-repress Dscam2.10A in R cells

